# Microprocessor dynamics shows co- and post-transcriptional processing of pri-miRNAs

**DOI:** 10.1101/097311

**Authors:** Annita Louloupi, Evgenia Ntini, Julia Liz, Ulf Andersson Ørom

**Author notes:** Correspondence to (U.A.Ø).

## Abstract

miRNAs are small regulatory RNAs involved in the regulation of translation of target transcripts. miRNA biogenesis is a multi-step process starting with the cleavage of the primary miRNA transcript in the nucleus by the Microprocessor complex. Endogenous processing of pri-miRNAs is challenging to study and the in vivo kinetics of this process is not known. Here, we present a method for determining the processing kinetics of pri-miRNAs within intact cells over time using a pulse-chase approach to obtain nascent RNA within a 1-hour window after labeling with bromouridine. We show, that pri-miRNAs exhibit different processing kinetics ranging from fast over intermediate to slow processing and provide evidence that pri-miRNA processing can occur both co-transcriptionally and post-transcriptionally.

## Introduction

microRNAs (miRNA) are small RNAs that mediate posttranscriptional regulation of gene expression (1). miRNAs are transcribed as primary transcripts as long as 30kb and processed by the Microprocessor complex in the nucleus (2). The Microprocessor complex is the minimal complex required for pri-miRNA processing in vitro consisting of the two proteins Drosha and DGCR8 (2).

Several features, such as sequence-motifs around the precursor miRNA (pre-miRNA) hairpin in the primary miRNA (pri-miRNA) transcript (3, 4) and within the hairpin loop (3) have been shown to be involved in processing efficiency in mammals. Furthermore, early studies of pri-miRNA transcripts have reported a co-transcripitonal processing by the Microprocessor complex (5, 6) using mostly in vitro assays and studies of single endogenous examples. The retention of pri-miRNAs at chromatin by factors such as HP1BP3 has also been suggested to be important for efficient pri-miRNA processing (7). The state-of-the-art methodologies are, however, not able to address the general and in vivo dynamics of pri-miRNA processing.

We have previously shown, that sequencing of the chromatin-associated RNA can reveal the steady-state processing efficiency of individual pri-miRNAs within the cell, demonstrating pri-miRNA processing as one of the most important factors for determining the level of mature miRNAs (4). To further follow processing of pri-miRNAs endogenously over time without constraints of their cellular localization or differential transcription rates we followed a nascent RNA sequencing protocol (8). We show that pri-miRNAs exhibit different processing kinetics both within the same polycistronic transcript and with respect to transcription and release from chromatin.

## Results

### Set-up of nascent RNA pulse-chase sequencing

RNA-sequencing yields an average view of RNA in the cell or in the respective purified subcellular compartment, reflecting a mixture of RNA of different age compared to the time of transcription. To follow RNA from transcription through processing, nascent RNA can be obtained by labeling actively transcribed RNA with a pulse of a modified nucleotide (*e.g.* Bromouridine (BrU) (8)) that allows for subsequent purification.

To expand our previous studies of steady-state pri-miRNA processing efficiency we used nascent RNA obtained after a short (15 min) BrU pulse and subsequent chase for 0, 15, 30 and 60 minutes (Samples 15, 30, 45 and 75 min after BrU, respectively) to follow the kinetics of the processing (Figure 1a). We subjected total nascent RNA from all time points obtained from HEK293 cells to next-generation sequencing using an Illumina Hi-Seq 2500 to obtain around 200 M reads per sample.

**Figure 1.**
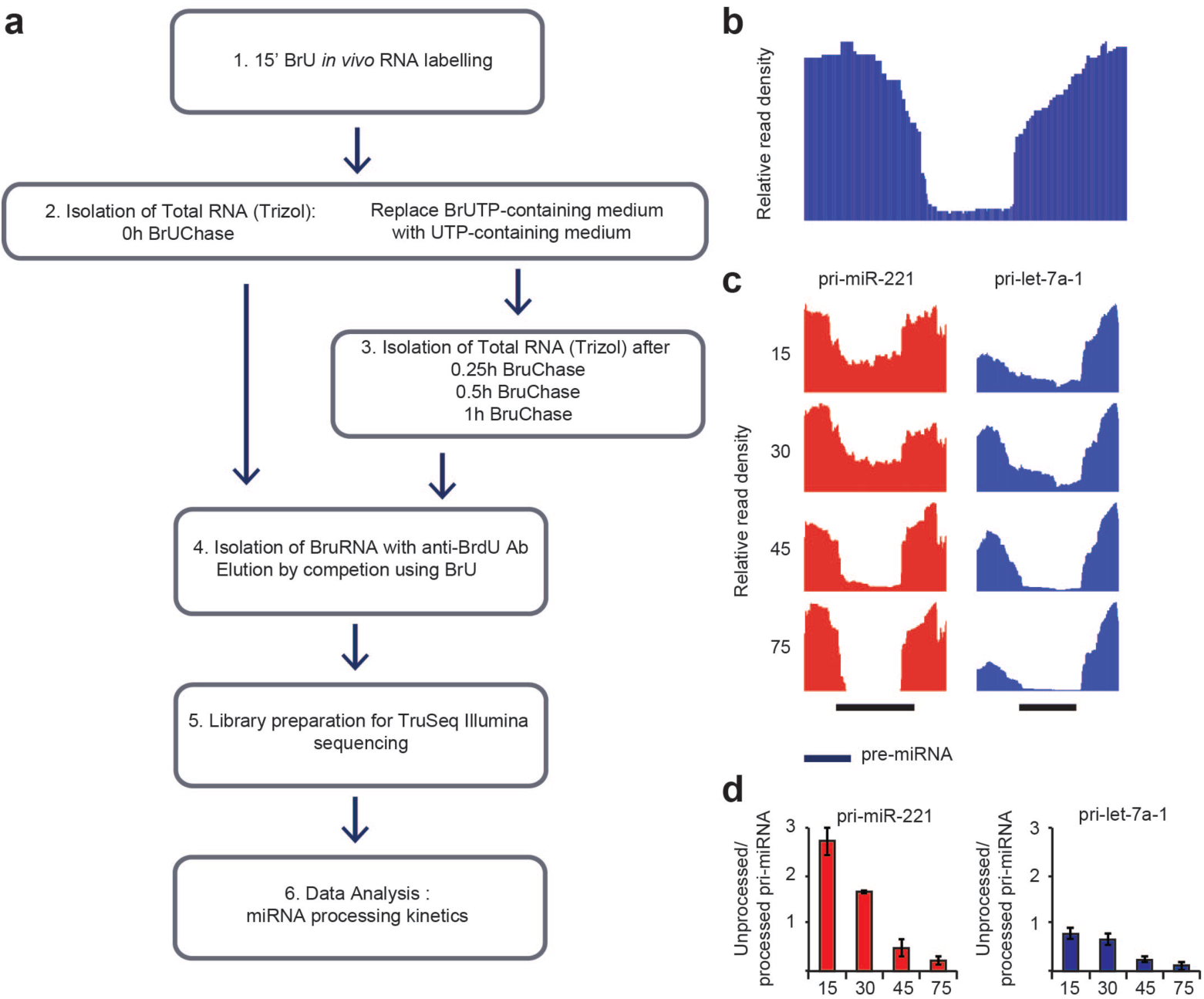
Measuring pri-miRNA processing kinetics with nascent RNA labeling. *a) Workflow for RNA pulse-labeling with BrU and chase to follow nascent RNA. b) Concept of processing signature in pri-miRNAs. Processing extent is calculated as the read-density in the pre-miRNA region compared to the flanking regions. Processing efficiency is calculated as (1 – processing extent). c) Processing signatures in RNA-sequencing data from nascent RNA in pulse-chase experiment for pri-miR-221 and pri-let-7a-1. d) Quantification by PCR of unprocessed/processed pri-miRNA for examples shown in c from two independent experiments.*

### In vivo profiles of processing kinetics from whole cells

We have previously reported a characteristic profile for steady-state chromatin-associated RNA around the site of pre-miRNA processing within the pri-miRNA transcript (4) (Figure 1b). Following the nascent RNA during the chase we can follow the time-course of processing of pri-miRNAs in HEK293 cells. Interestingly, for the 38 pri-miRNAs where we see a pronounced profile (4) (Supplementary Table 1) we observe different processing kinetics across pri-miRNA transcripts, and within polycistronic pri-miRNAs. The profiles for miR-221 and let-7a-1 are shown in Figure 1c, representing an intermediate processed (t1/2 ~ 40 min) and a fast processed (t1/2 < 15 min) pri-miRNA, respectively. These processing efficiencies can be recapitulated using quantitative PCR of individual pri-miRNAs with primers spanning the processing site to determine relative amounts of unprocessed pri-miRNAs, and primers amplifying the total of pri-miRNA transcript (processed + unprocessed), as described in (4). Representative results for miR-221 and let-7a-1 are shown in Figure 1d.

### Pri-miRNAs show differential kinetics in processing

These data show that some pri-miRNAs show almost complete processing at time point 0h, while others exhibit a slower pattern of processing by the Microprocessor (Supplementary Table 1). For several pri-miRNAs we observe a very low processing efficiency at 15’ (after 15’ BrU labeling, 0 min chase), arguing that we are able to capture some pri-miRNAs even before the processing by the Microprocessor complex takes place. Based on the processing kinetics we group pri-miRNAs into three groups: Fast processed (Figure 2a), Intermediate processed (Figure 2b) and Slow processed (Figure 2c). For Fast processed pri-miRNAs we observe the majority of Microprocessor activity within the pulse (before 15’), while for Intermediate processed pri-miRNAs the majority of processing happens around 30’-45’. For the group of Slow processed pri-miRNAs most processing occurs after 45’. While the fast and the intermediate processed pri-miRNAs all obtain a complete processing within 75’, several of the slow processed pri-miRNAs are less than 50% processed at the 75’ time point. The presence of sequence motifs around the pre-miRNA and within the hairpin-loop have been shown to affect processing efficiency of pri-miRNAs (3, 4), namely the UG(-14) 14 nts upstream of the pre-miRNA (3); GC(-13) 13 nts upstream of the pre-miRNA (4); the CNNC motif 16-18 (3) or 16-24 (4) nts downstream of the pre-miRNA as well as the GUG or UGU stem-loop motif (3). While these motifs are reported to associate with better processing only the CNNC motif is abundantly present in mammalian pri-miRNAs. In fact we see the CNNC motif in most pri-miRNAs analyzed with a preference for Fast over Intermediate over Slow processed pri-miRNAs, especially when considering the CNNC(16-24) motif (Figure 2d). We do not see a general enrichment of the UG(-14), GU(-13) or the GUG or UGU motifs for the analyzed pri-miRNAs (Figure 2d).

**Figure 2.**
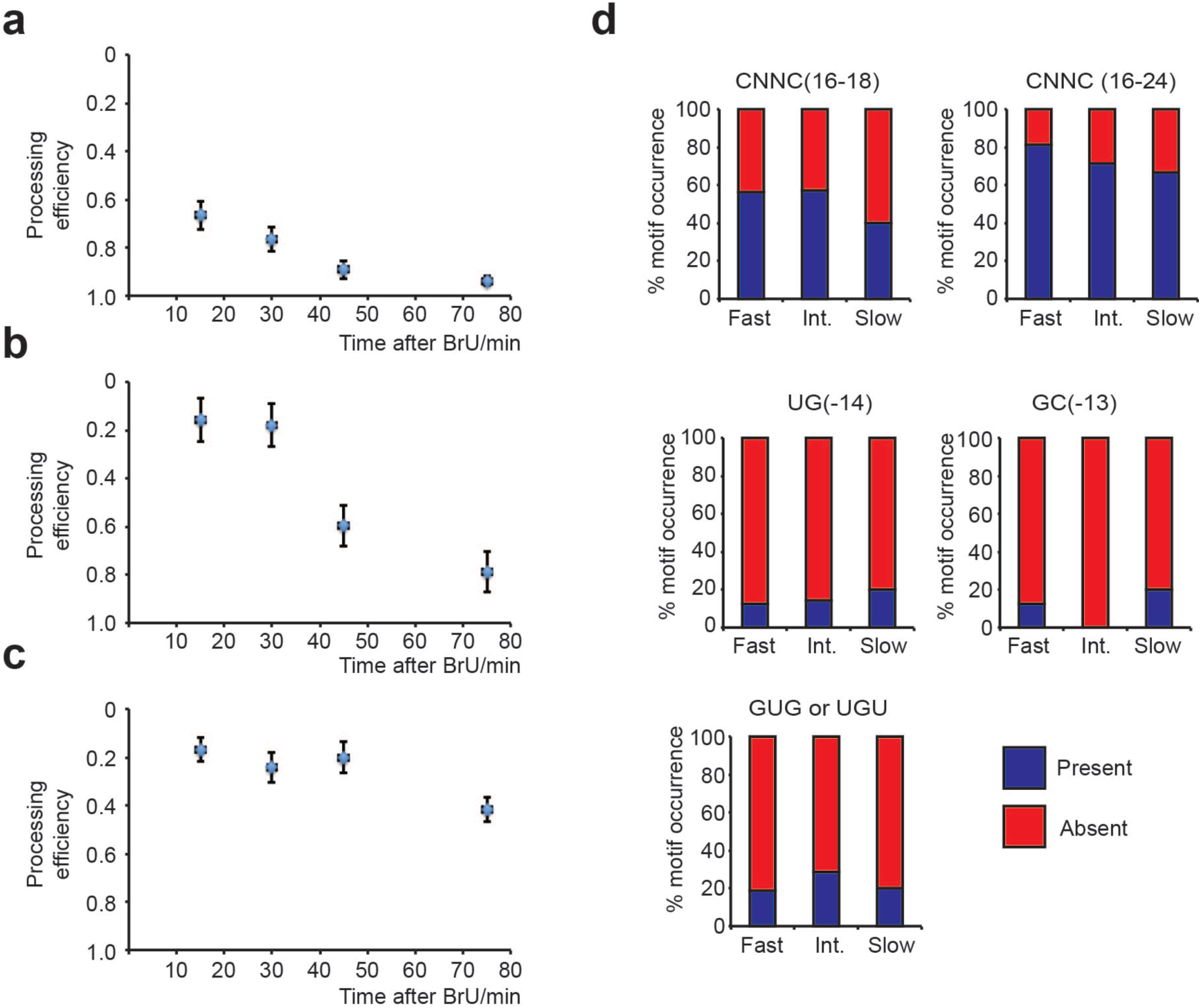
Pri-miRNA processing kinetics and associated motifs. *Average processing profile for a) Fast processed pri-miRNAs (n=16), b) Intermediate processed pri-miRNAs (n=7) and c) Slow processed pri-miRNAs (n=15). d) Motif occurrence for known motifs associated with pri-miRNA processing efficiency depicted as relative occurrence for each of the groups of pri-miRNA processing kinetics*.

### Differential processing within polycistronic pri-miRNAs

Several miRNAs are expressed from polycistronic pri-miRNAs (9) and these miRNAs often belong to the same families and thus predicted to target the same mRNAs for translation regulation and target RNA degradation (10). Prominent polycistronic pri-miRNAs are let-7a/f and miR-221/222, described to have important roles in cancer and cell cycle (11). We observe differential processing kinetics within both these polycistronic pri-miRNAs. The miR-221/222 pri-miRNA is a 25kb long transcript (Figure 3a) encoding two miRNAs. While the two miRNAs are relatively closely spaced, they exhibit very different processing kinetics (Figure 3b-c), demonstrating that processing kinetics are not defined by the primary transcript or its association to chromatin, as has recently been suggested (7). miR-221 and miR-222 are both shown to affect the cell cycle by targeting the p27 tumor suppressor and to promote cancer progression (12). The processing efficiency of miR-221 and miR-222 over time was quantified as described in Figure 1b and shown in Figure 3c. Interestingly, the expression levels of miR-221 and miR-222 in HEK293 cells are comparable and the half-life of the mature miRNAs have been estimated in mouse to be the same (13), suggesting that processing kinetics could define the biological regulation that could modulate the levels of miRNAs against a specific target through-out the cell cycle.

**Figure 3.**
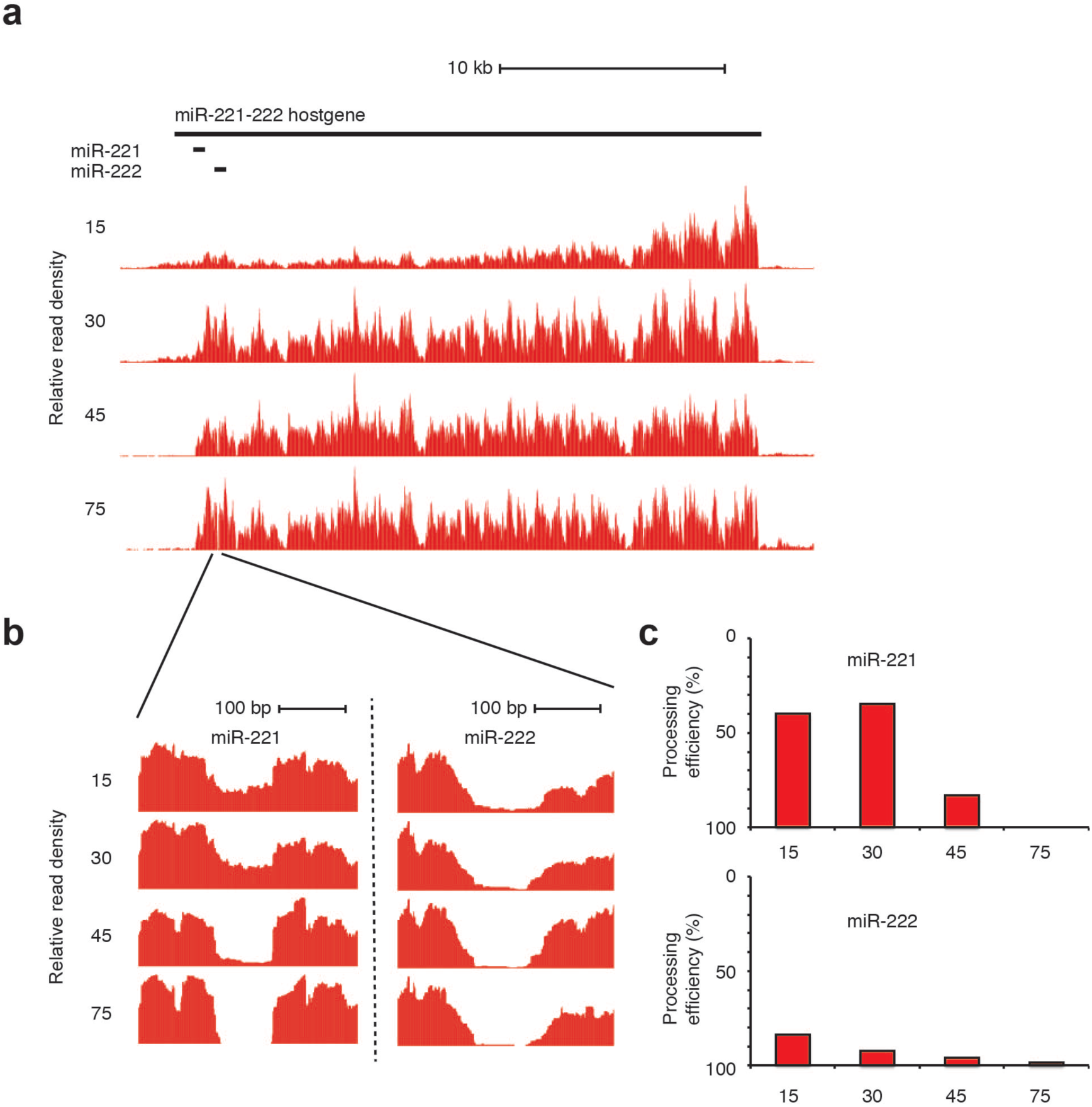
Differential processing within the pri-miR-221/222 polycistronic pri-miRNA transcript. *a) Overview of the genomic region and full pri-miRNA transcript. b) Enlarged read-densities around pre-miRNAs for miR-221 and miR-222. c) Quantification of processing efficiency from b.*

### Pri-miRNAs are processed co- and post-transcriptionally

The average processing efficiency as derived from steady-state chromatin-associated pri-miRNA data (4) should reflect the average processing efficiency of pri-miRNAs while associated to chromatin. The duration of an individual pri-miRNA association to chromatin prior release can be estimated by the dynamics of processing efficiency in time after pulse-chase of the nascent RNA, such that the average age of an individual pri-miRNA at chromatin is reflected by the corresponding processing efficiency. We used this assumption to estimate at what time transcripts are released from chromatin within each of the three groups of pri-miRNAs. Using the processing efficiency at chromatin derived from steady-state chromatin-associated RNA-sequencing data (4) and comparing it to the dynamics of the processing efficiency extracted for individual pri-miRNAs from the BrU pulse-chase sequencing data, we determine the half-life of pri-miRNA transcripts at chromatin (Figure 4a-c, shaded area). The Fast processed pri-miRNAs (Figure 4a) show very high processing efficiency when quantified from chromatin-associated RNA (4). While we estimate the chromatin-release of Fast processed pri-miRNAs to be 22-29 min (after beginning of pulse), the processing halflife is around 8 min (Figure 4a, dashed lines) arguing that these transcripts are truly co-transcripitonally processed and never leave chromatin. For the group of Intermediate processed pri-miRNAs we estimate chromatin-release to 34-48 min and the processing halflife to 41 min (Figure 4b, dashed lines). This suggests, that while co-transcriptional processing of this group of pri-miRNAs is generally inefficient the kinetics increase at the release of the pri-miRNA transcript from chromatin, or when the transcript is loosely associated to chromatin. The Very slow processed pri-miRNAs show little processing within the first 75 min and do not reach 50% on average within the 75 min applied in this study (Figure 4c).

**Figure 4.**
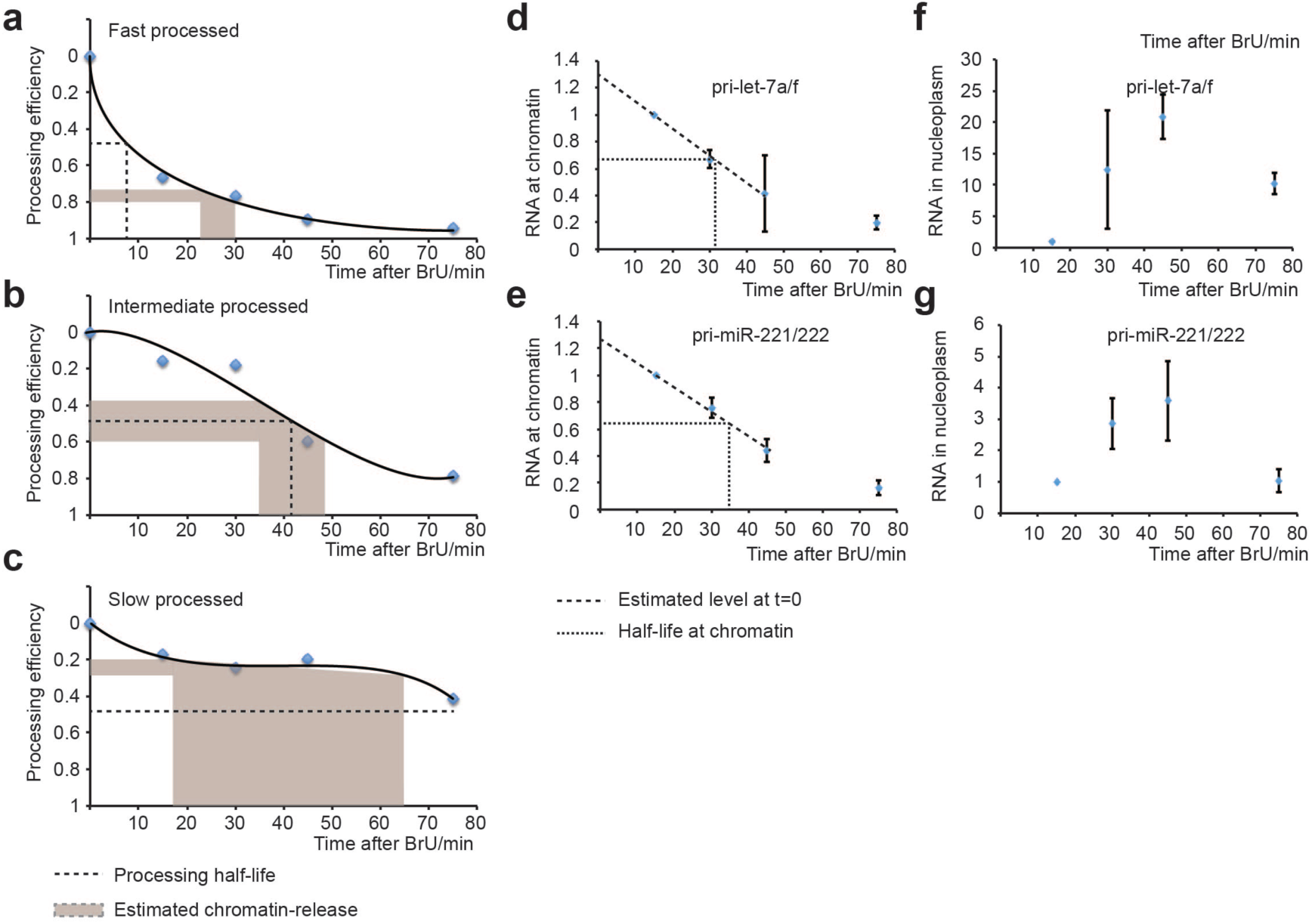
Chromatin-associated pri-miRNA processing and release. *Chromatin-release (shaded area) and processing half-life (dashed lines) are determined for a) Fast, b) Intermediate and c) Slow processed pri-miRNAs. d-e) Experimental validation of the release from chromatin of d) pri-let-7a/f and e) pri-miRNA-221/222. Dashed lines show estimated pri-miRNA at time 0’ (time of BrU addition) and fine dashed line show estimated half-life at chromatin. f-g) Relative amount of pri-miRNA in the nucleoplasm normalized to 15’. Experiments in d-g are from three independent experiments.*

To determine the dissociation rate of pri-miRNAs from chromatin, and to support our hypothesis that this can be estimated by the processing efficiency of chromatin-associated RNA, we isolated the chromatin and nucleoplasmic fractions of cells after a pulse labeling with BrU and chase for the same time-points (Figure 4d-g). Here, we show that for both miR-221/222 and let-7a/f the half-life at chromatin is 32-34 min, in agreement with the estimate from the dynamic pri-miRNA processing efficiency and in support of a model where Intermediate processed pri-miRNAs are more efficiently processed post-transcriptionally at their release from chromatin (Figure 4d-e). We furthermore show that pri-miRNAs accumulate in the nucleoplasmic fraction until 45’ after the pulse with BrU supporting an incomplete processing at chromatin (Figure 4f-g).

## Discussion

While previous work has proposed pri-miRNA processing to be an exclusively co-transcripitonal event (5, 6) it also suggested Drosha cleavage of the pri-miRNA as a relatively slow process (6). To this end we observe processing half-lifes of the groups of pri-miRNAs of 8 and 41 minutes respectively for the fast and intermediate processed pri-miRNAs. This is not in agreement with an exclusively co-transcriptional processing but fits well with Drosha cleavage being a relatively slow process. We observe, that the group of Intermediate processed pri-miRNAs are most efficiently processed at the time of release from chromatin or at a stage coinciding with the transcript half-life at chromatin where the transcript is being released from chromatin. This could be a state where the RNA becomes more available to catalytic nucleoplasmic proteins or less available to inhibitory proteins tightly bound to chromatin.

While factors responsible for chromatin-retention (7) or chromatin-release could play an important role in this process, we see chromatin-release of pri-miRNAs at comparable times after transcription for pri-miRNA transcripts showing different processing kinetics, arguing that chromatin retention is not the determining factor for pri-miRNA processing kinetics. In fact, the fast processed pri-miRNAs show a slightly faster release from chromatin than the intermediate and slow processed pri-miRNAs, in contrast to a model where a tighter association to chromatin increases processing efficiency as suggested in (7).

Drosha, the active part of the Microprocessor complex in cleaving pri-miRNAs, has been shown to be recruited to pri-miRNAs at chromatin co-transcripitonally (6), in some cases by the RNA-binding protein FUS (14). A possible scenario explaining that processing does not take place immediately for Intermediate and Slow processed pri-miRNAs could be that inhibitory factors associating to chromatin prevent co-transcriptional processing. As Drosha is recruited co-transcripitonally, these proteins could inhibit the activity of the chromatin-associated Microprocessor complex. Dissociation from chromatin would then lead to less interaction with these inhibitory factors and more efficient processing of the pri-miRNA transcripts. Identification of such factors would reveal important novel insight into the chromatin-RNA interactions and dynamics responsible for proper pri-miRNA processing and is an interesting challenge for future work.

## Materials and Methods

### Tissue culture and preparation of nascent RNA

HEK293 cells were cultured in DMEM growth-medium supplemented with 10% Fetal Bovine Serum (FBS) under normal growth conditions (37°C and 5% CO_2_). The day before bromouridine (BrU) labeling ~5.0 × 10^˄^6 cells were seeded in 150mm plates. Cells were 70-80% confluent before the addition bromouridine (BrU). BrU (-5-Bromouridine cat.no. CAS 957-75-5 Santa Cruz Biotechnology) was added to a final concentration of 2mM to the media and cells were incubated at normal growth conditions for 15 minutes (pulse). Cells were washed thrice in PBS and RNA purified by TRIzol after the chase time-points indicated. Labeled nascent RNA was purified using anti-BrdU antibodies conjugated to anti-mouse IgG magnetic Dynabeads. For elution 200μl of Elution buffer (0.1% BSA and 25mM bromouridine in PBS) were added directly on the beads and the tubes were incubated for 1h with continuous shaking (1100rpm) at 4 °C and RNA precipitated using ethanol.

For the chromatin-release assay cells were labeled with BrU and chased as described above. After the chase for the respective time points cells were fractionated as described in (15) on ice to obtain the chromatin-pellet and RNA extracted. Analysis using qPCR was done using the same number of cells for each condition.

### Quantitative real-time PCR

RNA was reverse transcribed using the High Capacity RNA-to-cDNA™ Kit (Invitrogen 4387406). cDNA was quantified on an 7900HT Fast real time PCR system (Applied Biosystems) using the SYBR^®^ Green PCR Master Mix (Invitrogen 4364344).

### RNA sequencing and data analysis

Library preparation was peformed using the Trueseq Stranded Total RNA Kit (Illumina). Sequening was performed on an Illumina HiSeq 2500 instrument to obtain around 200M reads per sample. Reads were mapped to GRCh37 (hg19) using STAR version 2.4.2.a (16) with default parameters. To assess the processing efficiency, we extracted the read counts covering the miRNA and the summed read counts from two surrounding 100nt intervals upstream and downstream of the 5’ and 3’ end of the miRNA respectively, after extending it by 20nt on both ends. The processing efficiency is the ratio of the miRNA read counts to the surrounding read counts, multiplied by the ratio 200 to the miRNA length.

### Filtering and annotation of miRNAs

microRNAs used in the analysis were filtered to include only high-confidence microRNAs showing absence of other non-coding RNA species in the region; folding of the pre-miRNA into a hairpin; and homogenous reads in small-RNA sequencing data for both the 5’ and 3’ mature miRNA. We required conservation of the hairpin structure in orthologous members of the gene family for conserved microRNAs (as defined in mirBase) including mouse or other mammals and conservation of the seed in more than 50 per cent of the orthologous genes. The miRNAs used is from (4) and includes 229 miRNAs; 138 classified as broadly conserved; 52 classified as weakly conserved; and 39 as non-conserved. We determined the exact Microprocessor cleavage sites using the annotation of the 5p and 3p miRNA strands from miRBase and mapped them onto the sequence of the pre-miRNA.

## Data access

Data from RNA sequencing are deposited in GEO with the accession number GSE…

## Author Contributions

A.L. did experimental work, interpreted data, conceived experiments and wrote the paper. E.N. did computational analysis, interpreted data and wrote the paper. J.L. did experimental work, conceived experiments and interpreted data. U.A.Ø. conceived the experiments, interpreted data, supervised research and wrote the paper. All authors read and approved the paper.

## Acknowledgments

E.N. has been funded by a postdoctoral stipend from the Alexander von Humboldt Foundation. Work in the author’s laboratory is funded by the German Research Council (DFG) and the Alexander von Humboldt Foundation through the Sofja Kovalevskaja Award to UAØ.

## Competing interests

The Authors declare no competing interests.

---

**Supplementary Table 1.** *Grouping and processing efficiencies ofpri-miRNAs included in the analysis.*

| Group | pri-miRNA | 15 min | 30 min | 45 min | 75 min |
| --- | --- | --- | --- | --- | --- |
| Fast | hsa-mir-9-2 | 0.96 | 0.13 | 0.06 | 0.00 |
|  | hsa-mir-301a | 0.62 | 0.12 | 0.11 | 0.00 |
|  | hsa-mir-30e | 0.58 | 0.17 | 0.14 | 0.00 |
|  | hsa-mir-218-1 | 0.44 | 0.81 | 0.46 | 0.07 |
|  | hsa-mir-10a | 0.39 | 0.21 | 0.10 | 0.05 |
|  | hsa-mir-423 | 0.32 | 0.16 | 0.02 | 0.00 |
|  | hsa-mir-30b | 0.31 | 0.17 | 0.02 | 0.01 |
|  | hsa-mir-98 | 0.34 | 0.11 | 0.03 | 0.11 |
|  | hsa-let-7a-1 | 0.24 | 0.22 | 0.06 | 0.02 |
|  | hsa-mir-101-1 | 0.22 | 0.49 | 0.03 | 0.03 |
|  | hsa-mir-222 | 0.16 | 0.08 | 0.04 | 0.02 |
|  | hsa-mir-103a-1 | 0.12 | 0.26 | 0.00 | 0.04 |
|  | hsa-mir-32 | 0.26 | 0.39 | 0.52 | 0.18 |
|  | hsa-let-7f-2 | 0.03 | 0.01 | 0.01 | 0.00 |
|  | hsa-mir-374a | 0.21 | 0.38 | 0.05 | 0.26 |
|  | hsa-mir-629 | 0.12 | 0.07 | 0.08 | 0.19 |
| Intermediate | hsa-mir-1307 | 1.18 | 0.91 | 0.23 | 0.08 |
|  | hsa-mir-505 | 0.92 | 1.05 | 0.49 | 0.61 |
|  | hsa-let-7d | 0.89 | 0.76 | 0.44 | 0.09 |
|  | hsa-mir-25 | 1.03 | 1.17 | 0.83 | 0.41 |
|  | hsa-mir-221 | 0.60 | 0.65 | 0.17 | 0.00 |
|  | hsa-let-7g | 0.75 | 0.74 | 0.43 | 0.20 |
|  | hsa-mir-378a | 0.52 | 0.47 | 0.24 | 0.09 |
| Slow | hsa-mir-616 | 0.88 | 0.78 | 0.82 | 0.35 |
|  | hsa-mir-590 | 0.93 | 0.85 | 1.14 | 0.42 |
|  | hsa-mir-641 | 0.93 | 0.66 | 0.91 | 0.58 |
|  | hsa-mir-197 | 0.83 | 0.82 | 0.75 | 0.51 |
|  | hsa-mir-545 | 0.90 | 0.70 | 0.79 | 0.60 |
|  | hsa-mir-186 | 1.13 | 0.88 | 0.92 | 0.32 |
|  | hsa-mir-573 | 1.04 | 1.04 | 1.13 | 0.77 |
|  | hsa-mir-491 | 0.75 | 0.94 | 0.73 | 0.49 |
|  | hsa-mir-7-1 | 0.58 | 0.65 | 0.39 | 0.39 |
|  | hsa-mir-561 | 0.92 | 0.57 | 0.58 | 0.74 |
|  | hsa-mir-550a-1 | 0.43 | 0.36 | 0.34 | 0.39 |
|  | hsa-mir-196a-1 | 0.97 | 1.34 | 1.16 | 0.95 |
|  | hsa-mir-1254-1 | 0.84 | 0.67 | 0.86 | 0.83 |
|  | hsa-mir-576 | 0.78 | 0.73 | 0.88 | 0.80 |
|  | hsa-mir-659 | 0.56 | 0.37 | 0.57 | 0.60 |

